# Mechano-inhibition of Endocytosis Sensitizes Cancer Cells to Fas-induced Apoptosis

**DOI:** 10.1101/2022.06.14.496195

**Authors:** Mehmet H. Kural, Umidahan Djakbarova, Bilal Cakir, Yoshiaki Tanaka, Yasaman Madraki, Emily T. Chan, Valeria Arteaga Muniz, Hong Qian, Jinkyu Park, Lorenzo R. Sewanan, In-Hyun Park, Laura E. Niklason, Comert Kural

**Author notes:** Present address: Humacyte Inc., Durham, NC, 27213, USA. Present address: Department of Medicine, Maisonneuve-Rosemont Hospital Research Center, University of Montreal, Montreal, QC, H1T 2M4, Canada. These authors contributed equally (Mehmet H. Kural, Umidahan Djakbarova).

## Abstract

The transmembrane death receptor Fas transduces apoptotic signals upon binding its ligand, FasL. Although Fas is highly expressed in cancer cells, insufficient cell surface Fas expression desensitizes cancer cells to Fas-induced apoptosis. Here, we show that the increase in Fas microaggregate formation on the plasma membrane in response to the inhibition of endocytosis sensitizes cancer cells to Fas-induced apoptosis. We used a clinically accessible Rho-kinase inhibitor, fasudil, that reduces endocytosis dynamics by increasing plasma membrane tension. In combination with exogenous soluble FasL (sFasL), fasudil promoted cancer cell apoptosis, but this collaborative effect was substantially weaker in nonmalignant cells. The combination of sFasL and fasudil prevented glioblastoma cell growth in embryonic stem cell-derived brain organoids and induced tumor regression in a xenograft mouse model. Our results demonstrate that sFasL has strong potential for apoptosis-directed cancer therapy when Fas microaggregate formation is augmented by mechano-inhibition of endocytosis.

---

Fas (CD95/APO-1) is a death receptor in the tumor necrosis factor (TNF) superfamily that regulates immune system homeostasis^1^. The physiologic ligand of Fas, FasL, is a type II transmembrane protein expressed on the surface of activated T cells and natural killer cells^2^. These cells can also secrete a soluble form of FasL (sFasL) that lacks the transmembrane domain and is therefore able to circulate freely in the bloodstream^3^. Upon ligation at the plasma membrane, Fas clusters form and generate the death-inducing signaling complex (DISC), which activates caspase-8 to initiate a downstream signaling cascade that leads to apoptosis^2,4^. Intracellular Fas trafficking, including recycling via endosomes to the plasma membrane, plays a vital role in ligand-induced DISC assembly^5,6^.

Since Fas expression is associated with increased cell proliferation and metastasis in numerous types of cancer, there have been several attempts to target Fas-mediated apoptosis to specifically kill tumor cells^7–15^. However, the development of a systemic therapy utilizing sFasL has thus far failed because sFasL alone has weak apoptosis-inducing capacity *in vivo*^8^ and most human tumors are desensitized to Fas-induced apoptosis^14,16^. Attempts to increase the efficacy of sFasL using cross-linking antibodies resulted in systemic toxicity in mice^8^. Thus, novel approaches to targeting the Fas-mediated apoptosis pathway are needed to take advantage of differences in Fas expression and signaling between cancer and normal cells.

The aggregation of Fas receptors in the plasma membrane increases the efficiency of DISC formation upon FasL stimulation^17,18^. As Fas levels below the threshold for ligand-induced clustering render cancer cells insensitive to Fas-induced apoptosis^6,19–21^, cancer cells have adopted distinct strategies to reduce Fas surface levels. For example, in tumor cells lacking functional p53, Fas is not transported from the Golgi complex to the plasma membrane^22^. Moreover, Fas molecules endocytosed from the cancer cell surface are predominantly delivered to lysosomes for degradation, not recycled to the plasma membrane^6^. Therefore, we aimed to determine whether slowing the endocytic machinery decreases Fas internalization and increases Fas density at the plasma membrane, ultimately increasing the susceptibility of cancer cells to Fas-induced apoptosis in the presence of sFasL through activation of the caspase cascade (Fig. 1a).

**Fig. 1:**
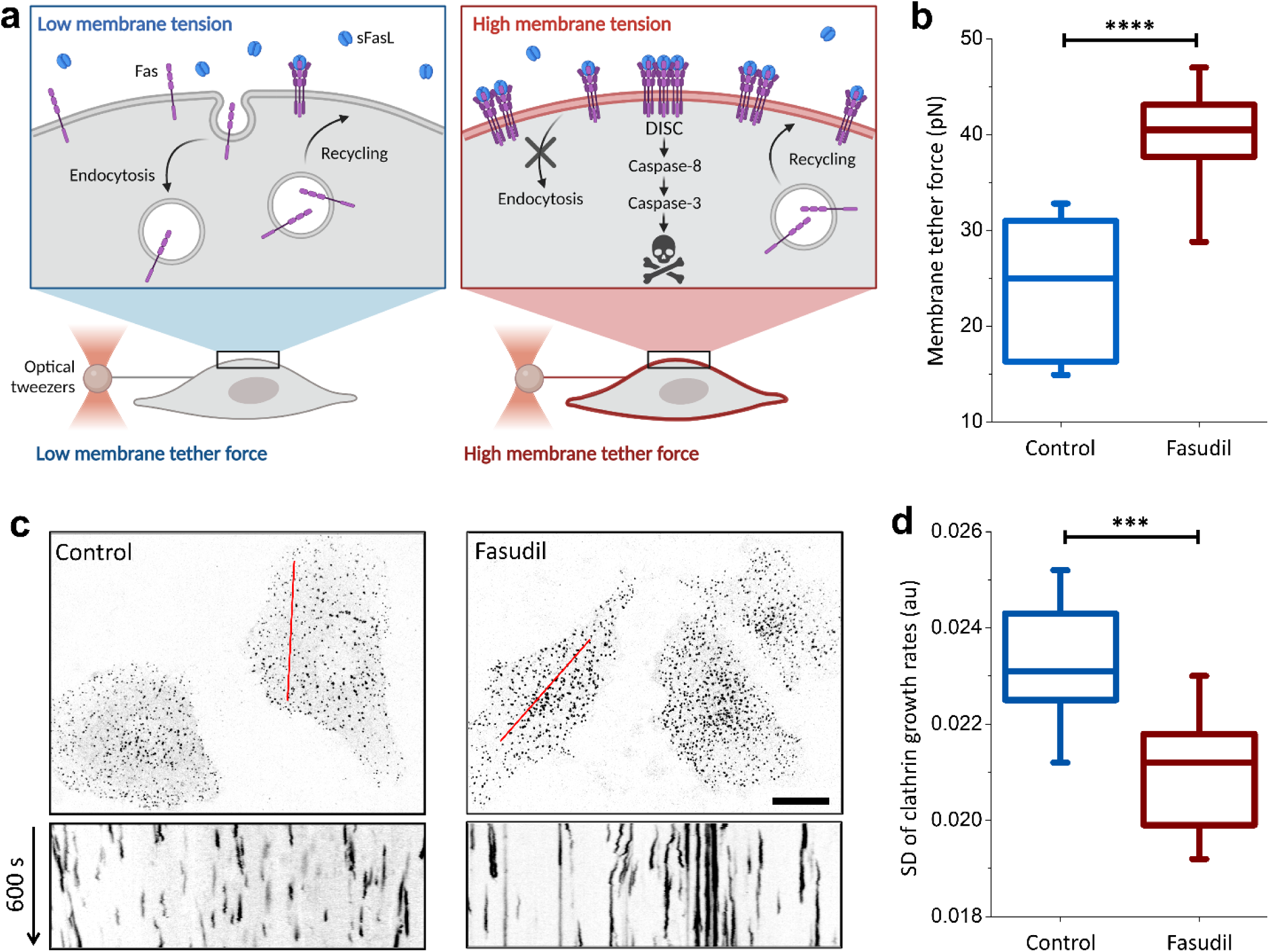
Mechano-inhibition of endocytosis by fasudil. **a**, Schematic representation of the study hypothesis: internalization of Fas from the cell surface reduces the sensitivity of cancer cells to Fas-induced apoptosis. Inhibition of endocytosis pathways is expected to increase Fas expression on the cell surface and enhance the formation of the death-inducing signaling complex (DISC) in the presence of sFasL. Endocytosis is slowed by increasing plasma membrane tension, which can be quantified using optical tweezers to measure the membrane tether force. **b**, SUM159 cells treated with 40 µM fasudil showed significantly higher membrane tension than did untreated cells. N_cells_ = 19 (untreated) and 28 (fasudil). **c**, Endocytic clathrin coats were imaged at the ventral surface of live SUM159 cells (genome edited to express AP2-EGFP) that were untreated (left) or treated with 40 µM fasudil for two hours (right). Kymographs obtained along the marked regions show clathrin-mediated endocytosis dynamics, with short streaks representing fast endocytic events. Streak length increased as endocytosis dynamics slowed. Fluorescence images are inverted to increase visibility. **d**, Standard deviation (SD) of the clathrin growth rates significantly reduced after treatment with 40 µM fasudil. N_cells_ = 14, N_events_ = 84230. ****p<0.0001, ***p<0.001; two-tailed t test. Scale bar, 20 µm.

Cells utilize endocytosis machinery to transform flat membrane patches into highly curved endocytic pockets and vesicles^23–26^; such curvature generation is hindered by mechanical tension on the plasma membrane, a potent and reversible regulator of endocytosis dynamics in cells^26–30^. Increasing membrane tension is one way to disrupt endocytosis and thus significantly alter the plasma membrane proteome^31,32^, thereby regulating various cellular and organismal processes^31,33–35^. Interestingly, the membrane tension of cancer cells is significantly lower than that of their nonmalignant counterparts^36–38^. Although lower membrane tension may set the stage for dysregulated endocytosis that contributes to malignant progression^39,40^, cancer cells are likely more susceptible to perturbations that impede endocytosis dynamics by elevating membrane tension^37,38^. In this study, we show that fasudil^41^, a clinically used Rho-kinase inhibitor, inhibits endocytosis in cancer cells by increasing plasma membrane tension. In support of our hypothesis, we found that inhibition of endocytosis increased the formation of Fas microaggregates on the cell surface and, thereby rendering cancer cells susceptible to Fas-induced apoptosis in the presence of sFasL. We further show that this strategy can be used to prevent glioblastoma growth in cortical organoids and mouse xenografts. Taken together, these results demonstrate that tumors can be targeted by Fas-induced apoptosis through the application of sFasL in combination with agents that reduce endocytosis dynamics and increase the formation of Fas microaggregates.

## Results

### Reduced actomyosin contractility by fasudil leads to mechano-inhibition of endocytosis

Mechano-inhibition of endocytosis is characterized in cultured cells using techniques that allow modulation of plasma membrane tension, such as cell stretching^27,42^, aspiration^43^, squeezing^28^, and hypo-osmotic swelling^28,44^. Although these approaches are effective in vitro, they are not suitable for systemic inhibition of endocytosis. Alternatively, reduced actomyosin contractility has been proposed to increase membrane tension by increasing the adhesion between the plasma membrane and the underlying actin cortex^45^. Correspondingly, we found that inhibition of the Rho-kinase by 40 µM fasudil treatment increased actin localization in the cell cortex (Supplementary Fig. 1a-d). The interaction between plasma membrane phospholipids and proteins and the underlying cortical actin is mediated by ezrin-radixin-moesin (ERM) family of proteins, known regulators of membrane tension^38,46,47^. Indeed, we found that fasudil treatment increased the peripheral localization of phosphorylated ERM along with actin (Supplementary Fig. 1e, f), and resulted in significantly higher membrane tension in SUM159 triple-negative breast cancer cells (Fig. 1a, b).

To assess the effects of fasudil on endocytosis, we focused on the dynamics of endocytic clathrin-coated carriers, as it was previously shown that the internalization of Fas is mediated by this pathway^48^. To this end, we used spinning-disk fluorescence microscopy to detect endocytic clathrin coats in live SUM159 cells that were genome-edited to express clathrin adaptor protein AP2 fused with GFP^49,50^. Using the AP2 signal as a proxy for clathrin coats allowed us to decouple endocytic activity from the clathrin-mediated membrane trafficking originating from intracellular organelles^51^. As expected, the endocytic clathrin coats on the adherent surface of SUM159 cells became increasingly static after fasudil treatment (Fig. 1c). To quantitatively assess the changes in endocytosis dynamics, we measured the standard deviation (SD) of the clathrin growth rates^43^, a robust indicator of clathrin-mediated endocytosis dynamics^28,29^, and observed a significant reduction following fasudil treatment (Fig. 1d). Contrary to fasudil treatment, increased actomyosin contractility by 30 nM leptin treatment for 24 hours resulted in significant reduction in membrane tension and, consequently, increased endocytosis dynamics (Supplementary Fig. 1c-e). Overall, our results demonstrate an inverse relationship between membrane tension and actomyosin contractility and that increased membrane tension by fasudil treatment mechanically slows endocytosis.

### Inhibition of endocytosis increases formation of Fas microaggregates on the cell surface

Given the ability of fasudil to slow endocytosis in cancer cells, we next aimed to investigate the effects of fasudil-mediated endocytosis inhibition on the membrane density and distribution of Fas receptors in a broad range of cancer cells. To accomplish this, Fas distribution in glioblastoma (U87), lung carcinoma (A549), liver carcinoma (HepG2), prostate cancer (PC3), and triple-negative breast cancer (SUM159) cells and in noncancerous human umbilical vein endothelial cells (HUVECs) was evaluated by immunofluorescence imaging. The Fas immunofluorescence signal was significantly higher in cancer cells than in HUVECs, in accordance with the previous studies reporting high levels of Fas expression in cancer cells^7–14^ (Fig. 2a). Fas receptors were observed as bright punctae on the surface of the cancer cells, indicating microaggregate formation (Fig. 2a; Supplementary Video 1). As the next step, we sought to examine the effect of fasudil-mediated inhibition of endocytosis by treating these cells with 40 µM fasudil for two hours. Upon treatment, the signal density of Fas microaggregates increased ∼2-fold on cancer cells, whereas no effect was observed on HUVECs (Fig. 2a-c). These results demonstrate that inhibition of endocytosis by fasudil treatment enhances the formation of Fas microaggregates on the surface of cancer cells.

**Fig. 2:**
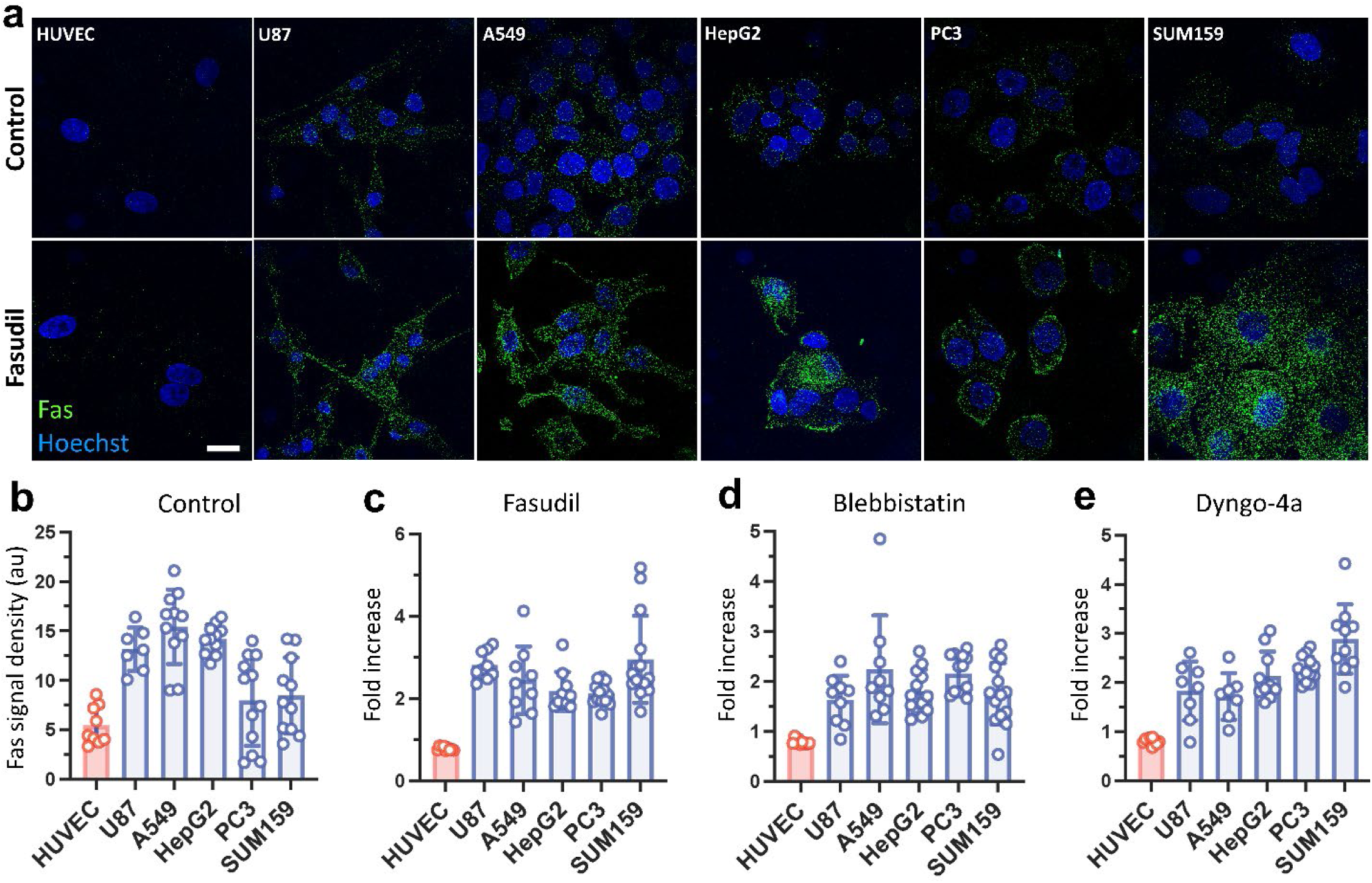
Inhibition of endocytosis increases Fas microaggregate density on the cell surface. **a**, Immunofluorescence assays show Fas microaggregate formation on HUVEC cells along with the cancerous U87, A549, HepG2, PC3 and SUM159 cells treated without (upper panel) or with fasudil (lower panel). **b**, The density of the Fas signal is quantified by measuring the integrated microaggregate fluorescence per cell area. Fas signal density is shown for different cells in the absence of endocytosis inhibition. **c-e**, Relative change in Fas signal density is shown for different cells after treatment with 40 µM fasudil (**c**), 10 µM blebbistatin (**d**), or 5 µM dyngo-4a (**e**) for 2 hours. Scale bar, 20 µm.

To corroborate the effects of endocytosis inhibition on Fas receptor density, we evaluated the formation of Fas microaggregates in cells treated with blebbistatin or dyngo-4a, which impede endocytosis dynamics mechanically and chemically, respectively. The myosin II inhibitor blebbistatin decreases actomyosin contractility in cells (Supplementary Fig. 1b), thereby increasing plasma membrane tension similar to fasudil^45,52^, whereas dyngo-4a inhibits receptor endocytosis by directly targeting the dynamin GTPase required for endocytic vesicle scission^53^. Consistent with the effects of fasudil, endocytosis inhibition by 10 µM blebbistatin or 5 µM dyngo-4a markedly increased the signal of Fas microaggregates on cancer cells treated for 2 hours, whereas no such increase was observed in HUVECs (Fig. 2d, e).

We next aimed to elucidate the mechanism behind the increase in signal density of Fas microaggregates upon endocytosis inhibition. Using flow cytometry to quantify protein expression, we found that 40 µM fasudil treatment for two hours results in no significant difference in the total expression levels of Fas (Supplementary Fig. 2a). To determine whether the increase in Fas signal density is exclusive to the cell surface, we used spinning disk confocal imaging to specifically analyze the ventral/adherent surface of the plasma membrane (Supplementary Fig. 2b). The fasudil– or dyngo-4a-mediated inhibition of endocytosis had no marked effect on the spatial density of Fas microaggregates on the adherent surface (Supplementary Fig. 2c). However, the mean immunofluorescence intensity of Fas microaggregates increased by up to one order of magnitude on the cancer cell surface but did not increase on the surface of noncancerous cells, i.e., HUVECs or human bronchial epithelial (HBE) cells (Supplementary Fig. 2d). Taken together, these results show that the inhibition of endocytosis does not affect the expression levels of Fas but promotes the growth of Fas microaggregates by increasing the number of Fas molecules on the cell surface.

### Inhibition of endocytosis sensitizes cancer cells to soluble Fas ligand

Given that the modulation of endocytosis increases Fas localization at the cell surface, we next sought to evaluate whether inhibiting endocytosis sensitizes cancer cells to exogenous FasL treatment. To this end, we assessed cell viability by quantifying ATP levels in culture medium after 48 hours of treatment with human recombinant sFasL alone or in combination with endocytosis inhibition. As shown in Fig. 3a, treatment with 400 ng/mL sFasL alone resulted in mild to moderate reduction in cancer cell viability (55.6%, 80.2%, 42.3%, 67.8%, 44.6%, and 59.9% in U87, PC3, BT549, A549, HepG2, and SUM159 cells, respectively, compared to the control). However, the addition of 40 µM fasudil increased the effect of sFasL, substantially decreasing the viability of the cancer cells (3.6-24.2%). As expected, unlike the effects observed in cancer cells, the viability of HUVECs, HBE cells, and induced pluripotent stem cell (iPS)-derived cardiomyocytes was minimally affected by the combination of fasudil and sFasL (84.0%, 75.9%, and 89.6%, respectively, compared to the control) (Fig. 3b; Supplementary Videos 2&3). Together, these results demonstrate that targeting endocytosis specifically increases the sensitivity of cancer cells to sFasL while sparing noncancerous cells.

**Fig. 3:**
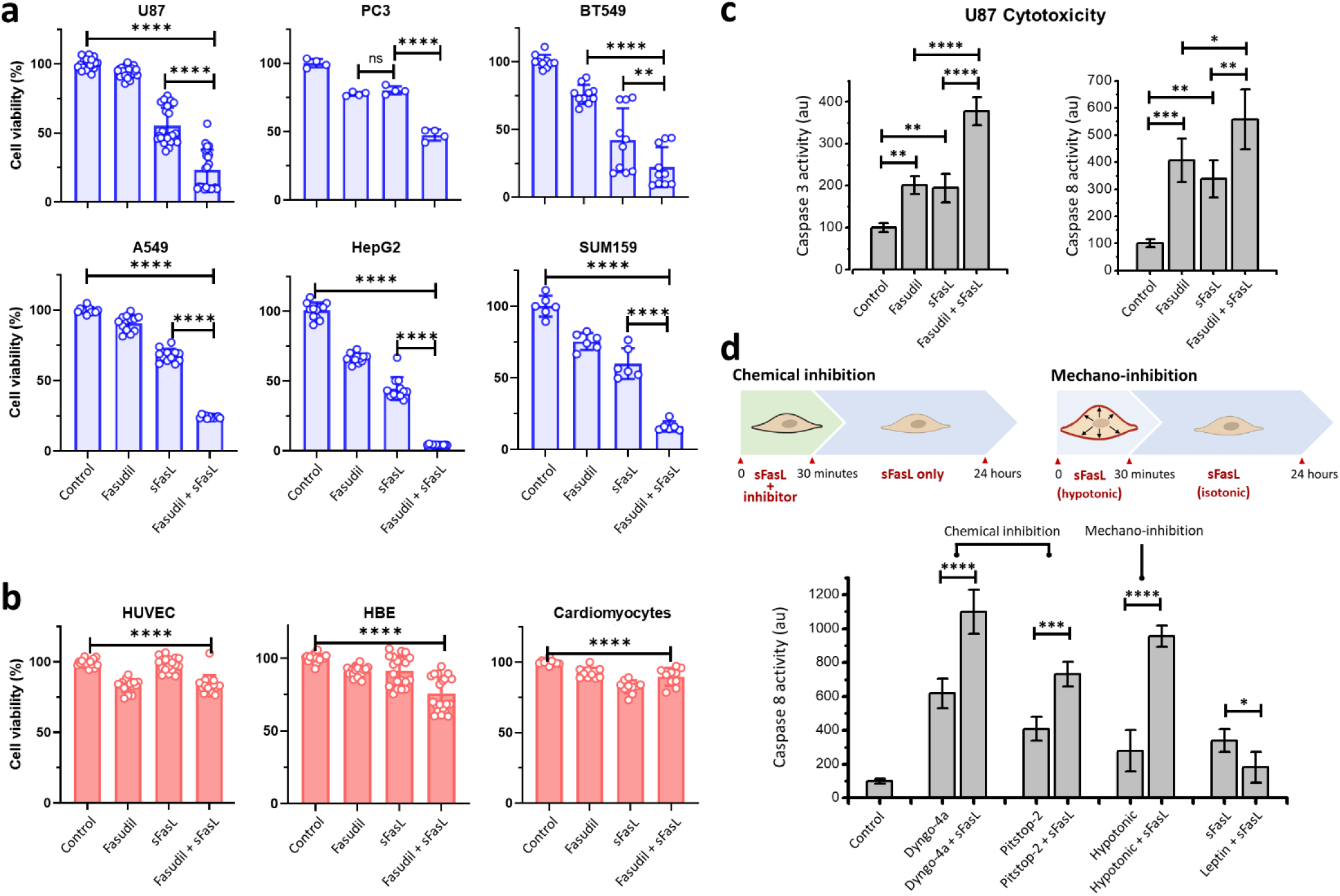
Inhibition of endocytosis sensitizes cancer cells to sFasL. **a**, Inhibition of endocytosis by 40 µM fasudil significantly decreased the viability of U87, PC3, BT549, A549, HepG2 and SUM159 cancer cells treated with 400 ng/mL sFasL. **b**, The viability of HUVECs, HBEs, and iPS-cardiomyocytes significantly decreased upon the same treatment, but this decrease was not as dramatic as that observed in cancer cells. **c**, Caspase-3 and caspase-8 activity in U87 cells significantly increased upon a combination of sFasL treatment with fasudil. **d**, Schematics describe the experiments where inhibition of endocytosis was limited to 30 minutes and caspase-8 activity was quantified after 24 hours (upper panel). Chemical inhibition was attained by treating cells with either 5 µM dyngo-4a or 2 µM pitstop-2. Mechano-inhibition involved increasing the plasma membrane tension of cells via hypotonic swelling for 30 minutes in the presence of sFasL. After this point, the hypotonic medium was replaced with isotonic medium (containing sFasL) to cease mechano-inhibition of endocytosis (****p<0.0001, ***p<0.001, **p<0.01; *p<0.05; one-way ANOVA).

To confirm that the reduction in viability is due to an increase in apoptosis, we monitored the activity of apoptotic markers caspase-3 and caspase-8 in U87 cells as we aimed to use these cells in our organoid and xenograft mouse models. In support of our hypothesis, we found that the inhibition of endocytosis upon fasudil treatment increased caspase-3 and caspase-8 activity in the presence of sFasL (Fig. 3c). To further corroborate that sensitization of U87 cells to sFasL is due to endocytosis inhibition, we performed cytotoxicity assays using alternative chemical and mechanical approaches to temporarily reduce endocytic rates. In our first approach, cells were pretreated with a combination of 400 ng/mL sFasL and a chemical endocytosis inhibitor (either 5 µM dyngo-4a or 2 µM pitstop-2) for 30 minutes and subsequently treated with 400 ng/mL sFasL only. After 24 hours, we found that even transient inhibition of endocytosis dynamics (for 30 minutes) significantly increased caspase-8 activity in the presence of sFasL (Fig. 3d). In our second approach, we relied on a purely mechanical strategy to slow endocytosis dynamics. Specifically, we temporarily increased membrane tension by incubating U87 cells with 400 ng/mL sFasL in 80% hypotonic culture medium for 30 minutes, as hypo-osmotic swelling during this period is known to result in increased membrane tension and slower endocytosis dynamics^28,30,54^. After 30 minutes, the mechano-inhibition of endocytosis was ceased by replacing the medium with isotonic medium (100% culture medium) containing 400 ng/mL sFasL. Quantification of caspase-8 activity after 24 hours revealed that a short exposure to hypotonic medium significantly increased the sensitivity of cancer cells to exogenous sFasL (Fig. 3d). On the contrary, caspase-8 activity in the presence sFasL reduced significantly when endocytosis dynamics are increased by 30 nM leptin treatment (Fig. 3d). To conclude, our findings demonstrate that inhibition of endocytosis increases the apoptotic efficacy of sFasL in cancer cells.

### Fas-mediated apoptosis targets glioblastoma in a brain organoid model

Next, we wanted to assess the potential utility of sensitizing cancer cells to Fas-induced apoptosis as a strategy for targeted cancer therapy by testing in physiologically relevant contexts. Therefore, we generated a 3D *in vitro* glioblastoma model based on embryonic stem cell-derived cortical brain organoids^55^ containing U87 glioblastoma cells labeled with red fluorescent protein (RFP). This approach allowed us to visualize and quantitatively assess changes in the glioblastoma mass within each organoid (Fig. 4a). Treatment with the combination of fasudil and sFasL (fasudil+sFasL) did not increase the percentage of apoptotic cells in 78-day-old brain organoids without U87 cells (Supplementary Fig. 3a, b). However, U87 cell-containing brain organoids treated with fasudil+sFasL for three days showed a dramatic decrease in the volume occupied by RFP-labeled U87 cells, leaving an acellular space (Fig 4b, c). The organoids stained positively for the neural progenitor marker SOX2 and the differentiated neural marker anti-β-Tubulin III (TUJ1) before and after the treatment (Fig. 4b; Supplementary Fig. 3c). Altogether, these results demonstrate that treatment with fasudil+sFasL specifically increases Fas-induced apoptosis in glioblastoma cells in brain organoids and has a minimal effect on the viability of neural progenitor cells.

**Fig. 4:**
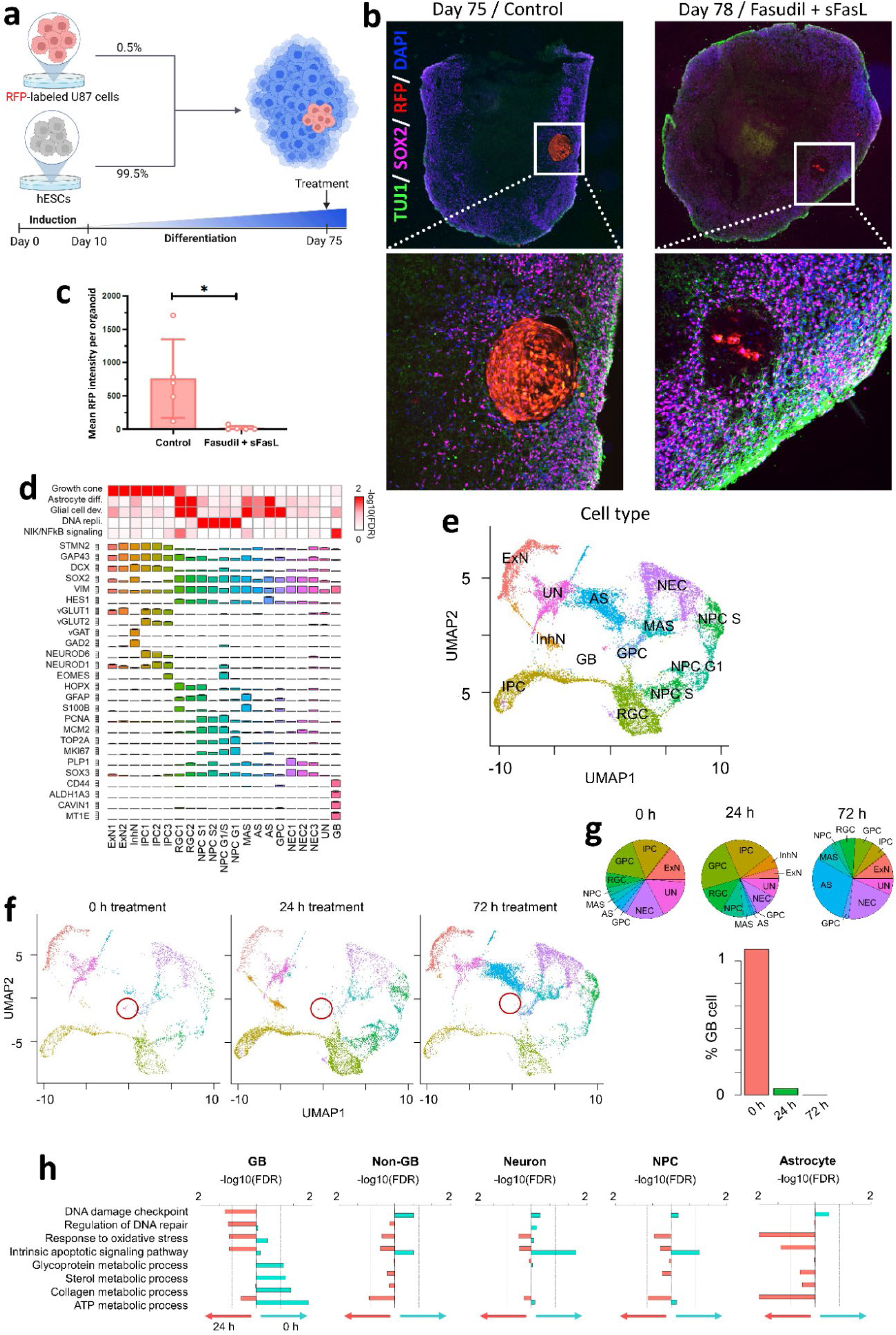
Fas-mediated apoptosis targets glioblastoma in a brain organoid model. **a**, Cortical brain organoids were formed by culturing human embryonic stem cells (hESCs, 99.5%) with red fluorescent protein (RFP)-expressing U87 glioblastoma cells (0.5%). **b-c**, Seventy-five-day-old organoids were imaged using spinning disk fluorescence microscopy, where the glioblastoma mass could be distinguished using RFP signal. Upon treatment with fasudil+sFasL, the RFP-expressing regions had almost disappeared by day 78 (p = 0.022; unpaired t test). **d-e**, Cell types in the u*niform manifold approximation and projection (*UMAP) plot of glioblastoma brain organoids. ExN: excitatory neurons, InhN: inhibitory neurons, IPC: intermediate progenitor cells, RGC: radial glia cells, NPC: neural progenitor cells, MAS: mature astrocytes, AS: astrocytes, GPC: glial progenitor cells, NEC: neuroepithelial cells, UN: unknown cell type, and GB: glioblastoma cells. **f**, UMAP plots of glioblastoma brain organoids after 0, 24, and 72 hours of treatment with fasudil+sFasL. The glioblastoma cluster (circled in red) shrank and eventually disappeared. **g**, Percentages of varying cell types in the organoid at the three different time points. The percentage of GB cells approached zero after 72 hours of treatment. **h**, After 24 hours of fasudil+sFasL treatment, the expression of genes related to apoptotic pathways, DNA damage, and oxidative stress increased in GB cells, but that of genes related to metabolic processes decreased.

To assess the effect of fasudil+sFasL on U87 glioblastoma cells in brain organoids at single-cell resolution, we performed single-cell transcriptome analysis with a total of 1391, 1097, and 2149 cells derived from organoids that were untreated, treated for 24 hours, or treated for 72 hours, respectively. We identified 21 clusters that were systematically classified into 13 cell types, including neurons, astrocytes, neural progenitor cells, and glioblastoma cells (Fig. 4d, e). Uniform manifold approximation and projection (UMAP) plots showed a dramatic decrease in the number of glioblastoma cells after 24 hours of treatment, with this population becoming negligible after 72 hours of treatment (Fig. 4f, g). A comparison of the global transcriptomes of untreated organoids and those treated with fasudil+sFasL identified 611 and 252 genes that were significantly up– and downregulated by treatment, respectively. Gene Ontology analysis revealed that the upregulated genes are enriched in the oxidative stress response (e.g., *GCLM*, *HSF1*, and *OGG1*), apoptosis (e.g., *MSH6*, *DNAJA1*, and *TRAP1*) and DNA repair (e.g., *FOXO4*, *MDM4*, and *CUL4A*) (false discovery rate (FDR) < 0.05 by hypergeometric test). In contrast, the glycoprotein (e.g., *MVD* and *OST4*), ATP (e.g., *COX5B* and *COX6C*), sterol (e.g., *NPC1* and *NPC2*) and collagen metabolic processes (e.g., *MMP2* and *VIM*) were significantly downregulated (FDR < 0.05) (Fig. 4h). The upregulation of cell death-related genes and downregulation of metabolic genes in response to fasudil+sFasL were more pronounced in glioblastoma cells than in noncancer cells, such as neurons, and neural progenitor cells. Interestingly, both apoptotic and metabolic genes were upregulated in astrocytes after treatment (Fig. 4h).

### Combination therapy of U87 glioblastoma in vivo xenografted tumors with fasudil and sFasL

Motivated by the targeted killing of U87 glioblastoma cells within cortical organoids, we further evaluated the antitumor efficacy of fasudil and sFasL as monotherapy and in combination using a subcutaneous U87 cell xenograft tumor model in athymic nude mice. In our preliminary in vivo assays, we found that, although high doses of sFasL result in systemic toxicity, the combination of 50 mg/kg Fasudil + 180 µg/kg sFasL is tolerated by mice upon intraperitoneal delivery (see Supplementary Fig. 4). However, this treatment led to no significant difference in tumor growth between the treatment arms (i.e., sFasL monotherapy, fasudil monotherapy and fasudil+sFasL) and the vehicle arm (Supplementary Fig. 4). Next, in an attempt to increase local drug concentration, the drugs were administered intratumorally twice a week in the beginning of the assay and the treatment route was switched to intraperitoneal administration after two weeks. We found that, 26 days after tumor cell implantation, the combination treatment (i.e., fasudil+sFasL) significantly inhibited tumor growth compared to vehicle or monotherapy (Fig. 5a). The median survival of mice treated with vehicle, fasudil or sFasL monotherapy ranged from 27-28 days, with fewer than 40% of the animals surviving one month after tumor cell implantation (Fig. 5b). Whereas, 26 days post implantation, while 37.5% (3/8) of mice in the control group reached endpoint, none of the mice in the combination treatment arm reached endpoint and tumor growth was observed in only one mouse. Fifty percent (4/8) of the mice in the combination arm showed tumor regression, and notably, one of the mice showed complete tumor regression (Fig. 5c). These results demonstrate that the local administration of fasudil+sFasL, but not either alone, significantly suppresses glioblastoma tumor growth, with potential curative effects *in vivo*. Overall, our results in cortical organoids and mouse xenograft tumor models demonstrate that sFasL can potentially be utilized as an effective and selective anticancer drug when combined with fasudil-mediated endocytosis inhibition.

**Fig. 5:**
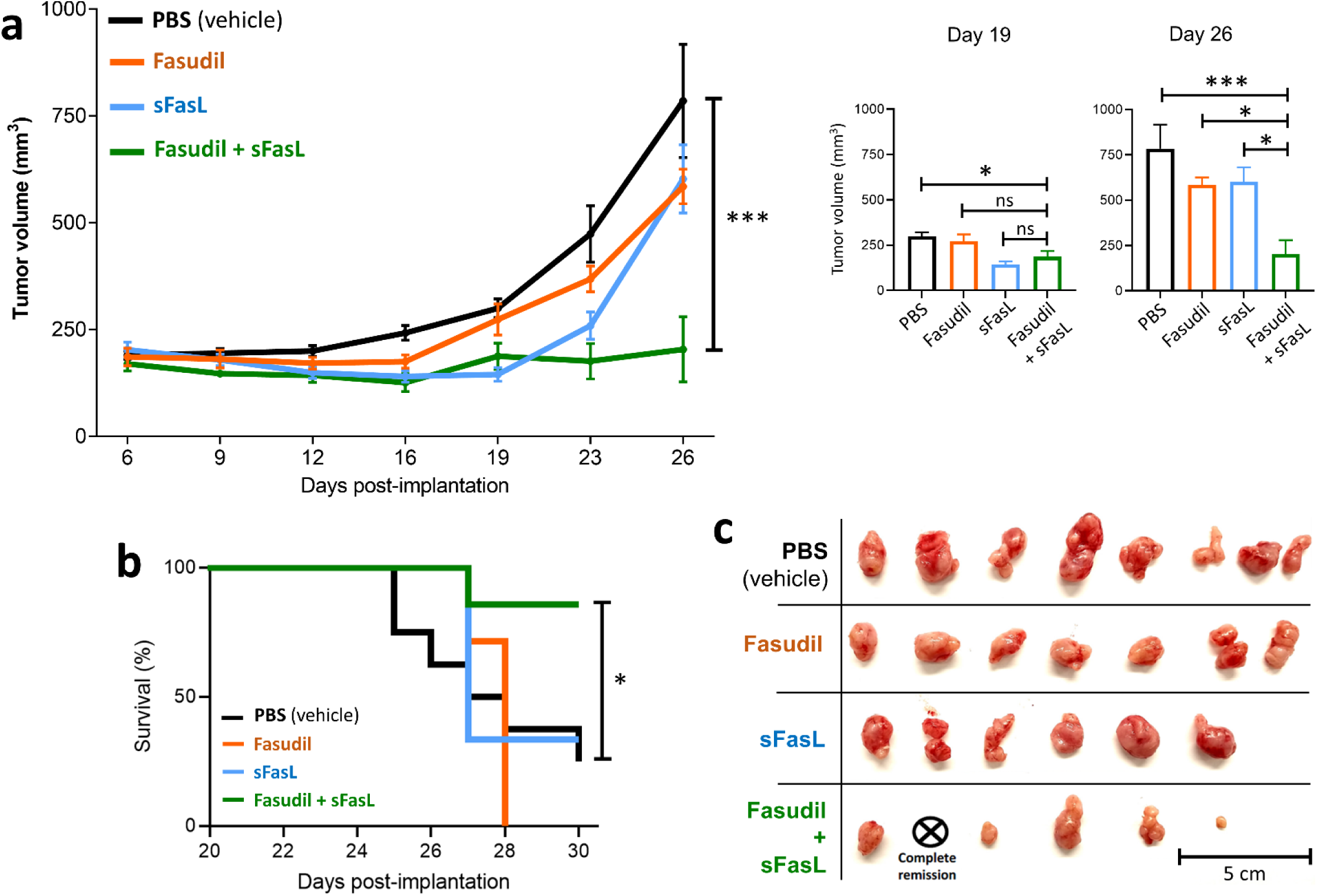
Local administration of fasudil and sFasL suppresses U87 glioblastoma growth in vivo. **a**, Tumor sizes in fasudil+sFasL treatment arm was significantly smaller than those in any other group at the end of the treatment. Panels on the right show the tumor sizes (mean ± sd) measured on 19– and 26-days post-implantation (***p<0.001, **p<0.01, and *p<0.05; ordinary one-way ANOVA). **b**, The survival probability in the fasudil+sFasL group was significantly higher than that in any other group [log-rank (Mantel‒Cox) test; compared to the combination group: PBS p=0.022, fasudil p=0.006, and sFasL p=0.048]. **c**, Extracted tumors one month after implantation. The tumor in one of the mice in the fasudil+sFasL combination group was completely eliminated.

## Discussion

Dysregulation of endocytosis has been linked to pathogenesis of numerous diseases^56–58^. Endocytosis dynamics is elevated in cancer cells to help reduce the expression of membrane proteins^56,59,60^, including death receptors^61,62^. Herein we show that fasudil, a clinically used vasodialator, hinders endocytosis dynamics in cancer cells by increasing the mechanical tension on the plasma membrane. Our results suggest that the increase in membrane tension is due to increased membrane-cytoskeleton adhesion upon reduced acto-myosin contractility, as proposed earlier^45^. Mechano-inhibition of endocytosis will enable further dissection of endocytosis function at the systemic level in different disease states, including cancer.

Cancer cells co-opt mechanisms to exploit Fas expression as a survival factor while circumventing Fas-induced apoptosis^15^. Our study aimed to sensitize cancer cells to Fas-induced apoptosis to transform this cell survival strategy into targeted cancer therapy. We show that inhibition of endocytosis promotes the formation of Fas microaggregates on the cell surface and thus decreases the ability of cancer cells to resist Fas-induced apoptosis. In a physiological context, this sensitivity can be activated via mechano-inhibition of endocytosis by fasudil. Overall, our results in cortical organoids and a mouse xenograft tumor model demonstrate that sFasL combined with fasudil has potential utility as an effective and selective anticancer therapeutic regimen.

Our results demonstrate that endocytosis inhibition does not affect the expression of Fas but increases the surface availability, promoting Fas microaggregation on the cancer cell surface. So far, there have been opposing reports regarding whether internalization of TNF family receptors form the cell surface is required for the induction of death receptor-induced apoptosis^48,63,64^. Our findings support models proposing that the internalization of the death receptors from the plasma membrane is not required for activation of the caspase cascade^64^.

We note that the cytotoxic activity of sFasL is orders of magnitude lower than that of membrane delivered FasL^8^. It was previously proposed that tumor cells release sFasL as a means to antagonize the effects of membrane-bound FasL^8,65^ or evade immune response^66^. Further investigations may establish whether sFasL shed by cancer cells may account for the increased caspase-8 activity observed in our assays upon endocytosis inhibition (mechanical or chemical) even in the absence of exogenous sFasL.

Finally, emerging research shows that cancer cells exhibit significantly lower membrane tension compared to their non-malignant counterparts^36–38^. Future investigations may delve into whether this reduced plasma membrane tension renders cancer cells more susceptible to mechano-inhibition of endocytosis. Moreover, exploring the mechanical characteristics of cancer cells could unveil opportunities to enhance the specificity of targeted therapies. This line of inquiry holds promise for advancing our understanding of cancer biology and optimizing therapeutic strategies.

## Materials and Methods

### hESCs culture

BC4 hESCs^55^ were cultured on Matrigel (BD Biosciences) coated dishes with mTeSR1 media (Stem Cell Technologies) and passaged every week by disassociating with 0.83 U/mL Dispase (Stem Cell Technologies). All experiments including hESCs were approved by Yale Embryonic Stem Cell Research Oversight (ESCRO).

### Cancer cell culture

U87, A549, and HepG2 cell lines were purchased from ATCC and were cultured in DMEM supplemented with 10% FBS and 1% pen/strep (Thermo Fischer). PC3 and BT549 cells were donated by Yale Center for Precision Cancer Modeling (YCPCM) and cultured in F-12 (Thermo Fischer) or RPMI (Thermo Fischer) media supplemented with 10% FBS and 1% pen/strep, respectively. SUM159 cells genome edited to express AP2-EGFP were donated by Tomas Kirchhausen’s group at Harvard Medical School ^50^. SUM159 cells were cultured in F-12 media supplemented with 5% FBS, 1% pen/strep, 1mg/ml hydrocortisone (Sigma Aldrich) and 10mg/ml insulin (Sigma Aldrich). Cells were kept at optimal density by passaging when needed. In viability experiments, cells were treated with different combinations of 400 ng/mL human recombinant sFasL (Biolegend), 40 μM Fasudil HCl (SellekChem), 10 μM Blebbistatin (Sigma), 5 μM Dyngo-4a (Cayman Chemical), or 2 µM Pitstop-2 (Abcam), hypotonic medium (80% of medium was replaced with water) in 96-well plates for 48 hours. Cell viability was quantified by detection of ATP content in the culture medium with CellTiter-Glo® Luminescent Cell Viability Assay (Promega) by following the manufacturer’s instructions. To assess the rate of apoptosis, caspase-3 and caspase-8 activities were measured using the EnzChek Caspase-3 Activity Assay Kit and the CaspGLOW™ Fluorescein Active Caspase-8 Staining Kit, respectively. The fluorescence generated by active caspase-3 and –8 formation was monitored at 25°C at excitation wavelengths (λ_ex_ = 485 nm) and emission wavelengths (λ_em_ = 535 nm) using an Infinite M1000 Pro plate reader (Tecan, Baldwin Park, CA, USA). Leptin (Sigma) concentration used in the caspase assays was 30 nM.

### Immunofluorescence microscopy and image analysis

Cells were fixed with ice cold 3.7% paraformaldehyde (Sigma-Aldrich), permeabilized with 0.1% Triton X-100 (Fisher Biotech, Hampton, NH, USA), and blocked with PBS supplemented with goat serum (2%, v/v) (Southern biotech) and 0.1% Tween-20 (Thermo Fisher). Cells were then incubated with FAS Mouse anti-Human (Clone: 4F8H6, Invitrogen) at 1:500 dilution and the appropriate fluorochrome-conjugated secondary antibody (Alexa Fluor 647 anti-mouse IgG1, Fischer Scientific) for Fas immunostaining. For labeling of actin filaments and pERM, cells were treated with Phalloidin AF405 (Thermo Fischer Scientific, 1:2000 dilution) and anti-pERM (Cell Signaling Technology #3726, 1:500) for 10-15 minutes at room temperature, following either the blocking step or secondary antibody labeling, depending on the experimental design. For nuclei staining, cells were incubated with Hoechst (1:2000) for 2 minutes at room temperature after secondary antibody labeling.

For quantitative analysis of cell surface-associated Fas expression, fluorescent images of the immunostained cells were collected using a Nikon TIE fluorescence microscope equipped with a CSU-W1 spinning disk unit (Yokogawa Electric Corporation), 100× objective lens (Nikon CFI Plan-Apochromat Lambda, NA 1.45), sCMOS camera (Prime 95B; Teledyne Photometrics), and 488– and 640-nm excitation lasers with 100 mW of nominal power. Images were acquired at a rate of 0.25–0.5 Hz with a laser exposure of 100 ms per frame. Image acquisition was performed using NIS Elements software.

A comparative analysis of Fas expression was conducted using Fiji software (https://imagej.net/Fiji/Downloads) by quantifying signal intensities obtained from confocal sections. To quantify the number, size, and area fraction of Fas puncta, the surface area of each cell was demarcated and subsequently masked to create binary images by using the ‘analyze particles’ function of the same software ^67^. After background subtraction, original image sections were multiplied by the mask obtained from the same field of view. For each cell, the mean intensity of spots was calculated by dividing the integrated intensity of the selected area by the number of detected spots.

Actin rearrangement and the cellular distribution of pERM were assessed through quantification of actin and pERM signal intensities in the peripheral regions, defined as the outermost 20% of the whole cell diameter, against central regions, encompassing 60% of the cell diameter.

### Membrane tether force measurements

Membrane tether force measurements were conducted at 37° C using the C-Trap optical tweezers system (LUMICKS, Amsterdam, The Netherlands). SUM159 cells gene-edited to express AP2–eGFP ^50^ were plated in a custom-designed flow chamber 15-24 hours prior to the experiments and incubated at 37°C and 5% CO_2_. The flow chamber was composed of a microscope cover glass (No 1.5, Fisher Scientific), a layer of parafilm tape as a separator, and a 1.0 mm thick microscope slide (Fisher Scientific). This selection of thicknesses for the cover glass, parafilm, and glass slide is designed for the flow channel to be compatible with the working distance of our C-Trap objective and condenser.

Tethers were pulled from the plasma membrane using streptavidin-coated polystyrene beads with a mean diameter of 1.87 μm (SVP-15, Spherotech). Right before the experiments, F-12 growth medium was replaced with phenol red-free L15 imaging medium (Thermo Fisher Scientific) supplemented with 10% FBS (containing the beads) and the flow channel inlet and outlet were immediately sealed. The imaging medium was prepared by diluting the bead suspension down to 0.01% w/v, spinning and resuspending the suspension once in PBS and then in phenol red-free L15 medium supplemented with 10% FBS.

The trap stiffness was calibrated using the thermal noise spectrum. After calibration, the trap stage was moved at a speed of 1μm/s until the bead contacted the cell. After 4-10 seconds, the stage was moved in the opposite direction at a speed of 5μm/s to attain a tether length of 10μm. The tether was then held stationary for up to a minute to measure the static tether force as described previously^68^.

### Imaging clathrin-mediated endocytosis dynamics in live cells

Fluorescence imaging was performed using a Nikon TIE fluorescence microscope equipped with a CSU-W1 spinning disk unit (Yokogawa Electric Corporation), a 100× objective lens (Nikon CFI Plan-Apochromat Lambda, NA 1.45), a sCMOS camera (Prime 95B; Teledyne Photometrics), and 488 nm excitation lasers with 100 mW of nominal power and temperature-controlled chamber. NIS Elements image acquisition software was used for acquisition of 2D time-lapse movies.

Endocytic clathrin dynamics were captured in SUM159 cells genome edited to express AP2-EGFP ^50^. 15-24 hours prior to imaging, cells were plated on a glass bottom dish (MatTek Life Sciences) and incubated at 37°C with 5% CO_2_. Cells were imaged in phenol red-free L15 medium (Thermo Fisher Scientific) supplemented with 10% FBS and 5% (v/v) penicillin/streptomycin at a rate of 0.25 Hz with laser exposure of 100 ms per frame. Two-dimensional time-lapse movies were acquired using NIS Elements software.

Individual clathrin-coated structures were captured and endocytic events were detected using TraCKer and cmeAnalysis softwares^43,51,69^. Clathrin coat growth rate analysis was conducted was previously described^43^ to assess the changes in clathrin-mediated endocytosis dynamics.

### Glioblastoma model in cortical brain organoids

As described previously^55^, hESCs were dissociated into single cells via Accutase and 9,000 cells, containing 0.5% RFP-infected U87 cells were plated into a well of U-bottom ultra-low-attachment 96-well plates. Neural induction medium (DMEM-F12, 15% (v/v) KSR, 5% (v/v) heat-inactivated FBS (Life Technologies), 1% (v/v) Glutamax, 1% (v/v) MEM-NEAA, 100 μM β-mercaptoethanol, 10 μM SB-431542, 100 nM LDN-193189, 2 μM XAV-939 and 50 μM Y27632) was used as culture media for ten days. Organoids were transferred to ultra-low-attachment six-well plate in hCO media without vitamin A (1:1 mixture of DMEM-F12 and Neurobasal media, 0.5% (v/v) N2 supplement, 1% (v/v) B27 supplement without vitamin A, 0.5% (v/v) MEM-NEAA, 1% (v/v) Glutamax, 50 μM β-mercaptoethanol, 1% (v/v) penicillin/streptomycin and 0.025% insulin) at day ten and the media was changed every other day. At day 18, Vitamin A, brain-derived neurotrophic factor, and ascorbic acid were added to the culture medium. Organoids with or without glioblastoma were treated with sFasL+fasudil combination after Day 75 for 72 hours for viability studies.

### Immunofluorescence microscopy and image analysis for organoids

Organoids were fixed in 3.7% paraformaldehyde at 4°C overnight followed by three washes with PBS at room temperature. Then, organoids were incubated in a 30% sucrose solution for 2 days at 4 °C. Organoids were equilibrated with optical cutting temperature compound at room temperature for 15 minutes, transferred to base molds, and embedded in optical cutting temperature compound on dry ice. Then, 40-μm cryosections were generated, and organoid blocks were stored at −80°C. Slides were dried for 2 hours at room temperature and incubated with 0.1% Triton-100 for 15 minutes at RT in a humidified chamber. Organoids were blocked with 3% BSA at room temperature for 2 hours, and then were incubated with the primary antibodies (mouse anti-Sox-2 and rabbit anti-TUJ-1) diluted 1:100 in 3% BSA overnight at 4°C. After two washing steps, organoids were incubated with Alexa Fluor Dyes (1:500) for 1 hour following nuclei staining with DAPI (1:1,000). Finally, slides were mounted with ProLong Gold Antifade Reagent and images were taken with a CSU-W1 spinning disk unit (Yokogawa Electric Corporation), 20× objective lens (CFI Plan Apochromat Lambda 20x/0.75, WD 1 mm, No: MRD00205), Andor iXon camera (Ultra888 EMCCD, 1024×1024 (pix), 13 um pixel), and 488-, 550-, and 640-nm excitation lasers with 100 mW of nominal power. DNA strand breaks were detected with TUNEL stain (11684795910, Sigma) to detect apoptotic or dead cells following the manufacturer’s protocol. The ratio of the mean signal intensity for TUNEL stain to DAPI was calculated for each image.

For quantitative analysis of TUJ1 and SOX2 staining in the organoids, manual masks of the maximum intensity projection of the images were generated using ImageJ. The DAPI signal in these masks was used to segment the nuclei and postprocess the data using the Python package scikit-image. Nuclei segmentation was performed using Cellpose, with a user-trained model. The masks retrieved by Cellpose were then expanded by three pixels radially using the function expand_labels (from sci-kit image) to encapsulate the entire cell. For each of the cell masks, the mean intensity in the TUJ1, SOX2 and DAPI channels was calculated using the function skimage.measure.regionprops and the background intensity was subtracted.

### Flow cytometry analysis

Cells were cultured under specified conditions, then washed with Phosphate-Buffered Saline (PBS) and detached using trypsin in preparation for counting and staining procedures. To evaluate Fas (CD95) expression, 3.0 × 10^5^ cells were suspended in 3.7% Paraformaldehyde (PFA) and incubated for 15 minutes at 4°C in the dark. This was followed by a permeabilization step with 0.1% Triton X-100 for 20 minutes at 4°C in the dark. After each step, cells were washed with PBS to remove any residual reagents. For blocking, cells were treated with 3% Bovine Serum Albumin (BSA) in Flow Cytometry Staining buffer (Thermo Fisher) overnight (ON). Subsequently, cells were stained with either CD95 APC antibody (Clone 1507, Thermo Fisher) or the APC IgG1 antibody (Thermo Fisher) as an isotype control. The staining was performed for 45 minutes at 4°C in the dark. Following staining, cells were washed twice with PBS and kept on ice prior to flow cytometry analysis.

The stained cells were analyzed using a Becton Dickinson LSR II flow cytometer at the Ohio State University Flow Cytometry Shared Resources. Data analysis was conducted using FlowJo software (version 8.8.6, Treestar Inc). Initial gating with FSC-A and SSC-A was performed to exclude dead cells, and subsequently, FSC-W and SSC-W gating was employed to eliminate doublets. For each sample, ∼10,000 events were captured, and the mean fluorescence intensity of the CD95 APC was analyzed using FlowJo software.

### Data processing of single-cell RNA-seq

Single-cell RNA-seq reads were mapped to GRCh38 human genome (GRCh38-2020-A) and counted with Ensembl genes using the count function of CellRanger software (v3.0.2) with default parameters. Batch effect and intrinsic technical effects were normalized by Seurat (v3.0.2)^70^. Briefly, we filtered out cells with less than 50 detected genes and more than 15% of mitochondria-derived reads as low-quality samples. Raw UMI count was normalized to total UMI count in each library. Top 2,000 highly variable genes were used to identify cell pairs anchoring different scRNA-seq libraries using 20 dimensions of canonical correlation analysis. All scRNA-seq libraires used in this study were integrated into a shared space using the anchor cells. After scaling gene expression values across all integrated cells, we performed dimensional reduction using principal component analysis (PCA). For the visualization, we further projected single cells into two-dimensional UMAP space from the 1^st^ and 20^th^ PCs. Graph-based clustering was then implemented with shared nearest neighbor method from the 1^st^ and 20^th^ PCs and 0.8 resolution value. Differentially expressed genes (DEGs) in each cluster were identified with more than 1.25-fold change and p<0.05 by a two-sided unpaired T test. Gene Ontology analysis was performed to the DEGs by GOstats Bioconductor package (v2.46.0)^71^. False discovery rate was adjusted by p.adjust function in R with “method=”BH”” parameter.

Cluster labeling was performed by unique markers, Gene Ontology, and enrichment of gene signatures as described previously^55,72^. Briefly, neuronal and non-neuronal clusters were separated by neuronal growth cone (*STMN2*, *GAP43*, and *DCX*) and early neurogenesis markers (*VIM*, *SOX2* and *HES1*). Neuronal clusters are further classified into excitatory and inhibitory neurons by expression of glutamate (*vGLUT1/2*) and GABA transporters (*vGAT*), respectively. In addition, neurons expressing neuroblast markers (*NEUROD1/6*) were labeled as intermediate progenitor cells (IPCs), which are transitioning to newborn neurons^73^. Non-neuronal clusters are classified into mitotic and non-mitotic cells with the presence and absence of cell cycle-related gene expression, respectively. The mitotic clusters were labeled as NPC S, NPC S/G1, or NPC G1 by S (*PCNA*, *MCM2*) and G1 phase markers (*TOP2A*, and *MKI67*). The non-mitotic clusters with significant overrepresentation of a GO term “Astrocyte differentiation (GO:0048708)” were assigned to astrocytes. Astrocyte clusters with high expression of *GFAP* and *S100B* were further labeled as mature astrocytes. *HOPX*-expressing non-mitotic clusters were assigned to radial glia cells. A non-mitotic cluster without astrocyte marker expression, but with significant overrepresentation of “glial cell development (GO:0021782)” was labeled as a glial progenitor cell. Neuroepithelial cells were assigned by expression of its markers (*PLP1* and *SOX3*). Glioblastoma clusters were assigned by unique expression of GBM biomarkers (*CD44*, *ALDH1A3*, *CAVIN1*, and *MT1E*). Global comparison of transcriptome profiles was performed by 1.25-fold average expression difference and p<0.05 of two-sided unpaired T test between 72h-treated and non-treated organoids.

### Fasudil+sFasL combination therapy in U87 glioblastoma xenograft tumors

Ten million U87 glioblastoma cells (ATCC) were implanted subcutaneously into the right flank of immune deficient athymic nude mice (Jackson Labs) in a 1:1 mix of plain growth media and Matrigel (Corning). After approximately one week post transplantation when the tumors were palpable, with tumor volumes ranging from 50-200 mm^3^ 7-8 tumor bearing mice were randomly assigned to the following four treatment arms: Vehicle (PBS), sFasL monotherapy, fasudil monotherapy and fasudil+sFasL combination. On Mondays and Thursdays, the Fasudil monotherapy and the fasudil+sFasL combination arms received a priming dose of 50 mg/kg Fasudil, while the sFasL only and PBS groups received an equivalent volume of PBS injection. Twenty-four hours after the priming injections, PBS, sFasL (180 μg/kg), fasudil (50 mg/kg) or fasudil (50 mg/kg) and sFasL (180 μg/kg) in combination, was administered to the relevant arms. Our maximum tolerated dose (MTD) studies demonstrated that the drug combinations used in this study do not cause fatality or significant weight loss in mice (Supplementary Fig. 4a).

In the first study, the route of administration was intra-peritoneal for the entire study. In the second study, the route of drug administration was intra-tumoral to the central tumor mass for two weeks but switched to intra-peritoneal due to observation of trauma and tissue dehiscence at the injection sites. Tumor dimensions were recorded by caliper measurements at three-day intervals and volumes calculated using the formula: (length x width^2^)/2. Treatment was ceased and the mice were euthanized after endpoint volume of 1000 mm^3^ was reached in the vehicle arm. Tumors were excised from mice from all arms and images taken at the end of the study.

## Acknowledgements

This work was funded by American Heart Association Postdoctoral Fellowship (17POST33661238) awarded to M.H.K., Pelotonia Fellowship to U.D. and E.T.C., NSF Faculty Early Career Development Program (award number: 1751113) and NIH R01GM127526 to C.K., and NIH R01HL148819 to L.E.N. Any opinions, findings, and conclusions expressed in this material are those of the authors and do not necessarily reflect those of the Pelotonia Fellowship Program or the Ohio State University. We thank Yale Center for Precision Cancer Modeling, for their help with the animal studies, and Yale West Campus Imaging Core. We also thank Dr. Dmitri Kudryashov, Dr. Elena Kudryashova and Junyan Yu from the The Ohio State University for helping us with the viability and apoptosis assays.

## Author Contributions

M.H.K. oversaw the project, designed, and optimized the experiments and analyzed the data. C.K and M.H.K. prepared the manuscript. U.D. performed the immunofluorescence experiments and analyses in cultured cells, and conducted the viability and cytotoxicity assays involving endocytosis inhibitors. U.D. and V.A. performed the flow cytometri assays. B.C. helped with the organoid experiments. V.A. analyzed the organoid images. Y.T. analyzed the single-cell RNA sequence data, Y.M. and E.Y.T. performed the tether force experiments. H.Q. helped with RFP labeling, J.P. and L.R.S. helped with iPS-derived cardiomyocytes. I.P. oversaw the organoid experiments, L.E.N. and C.K. oversaw the project.

## Supplementary Information

**Supplementary Figure 1.**
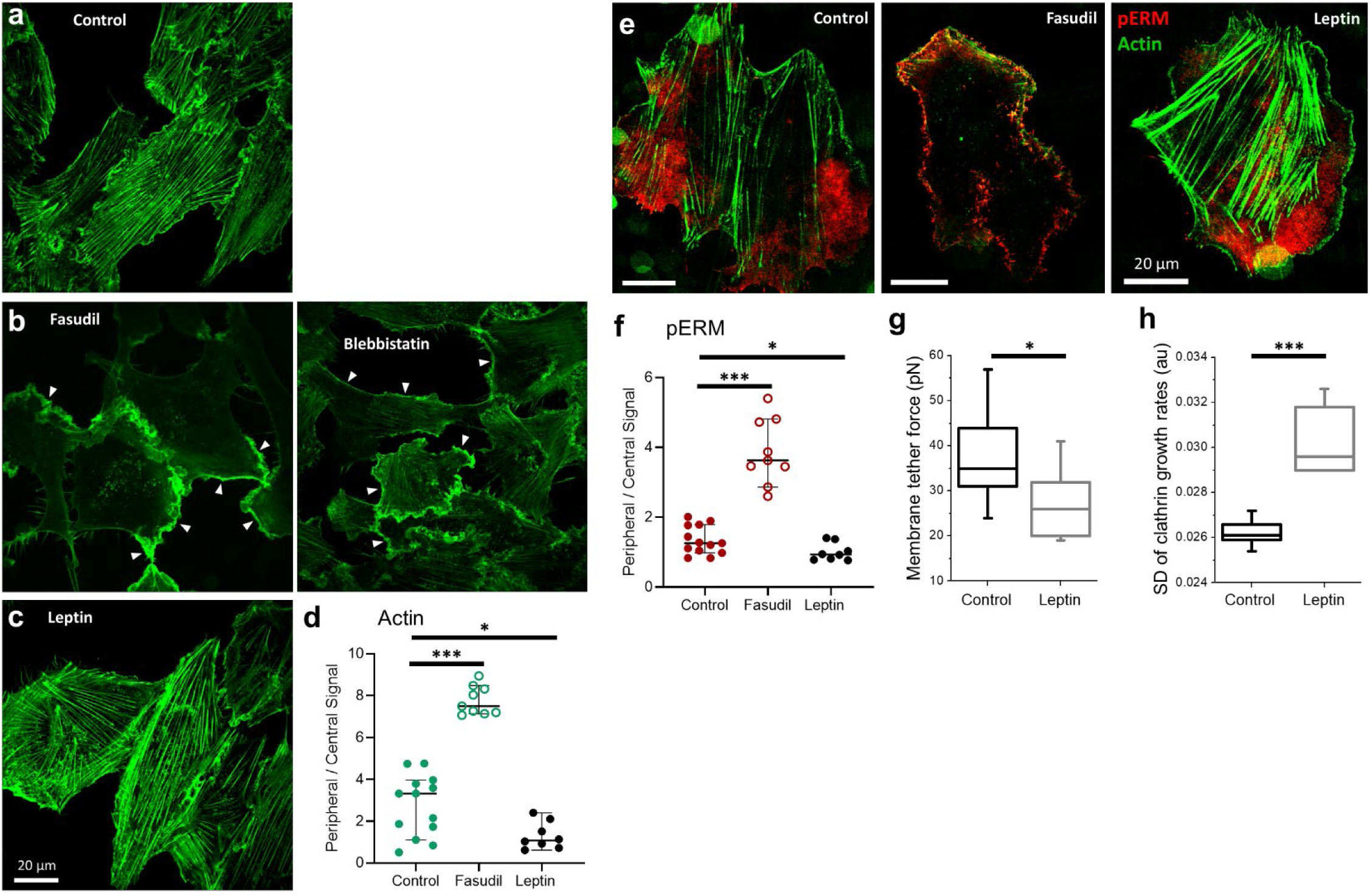
**a-c**, Immunofluorescence microscopy shows the distribution of actin in SUM159 cells under different conditions. **b**, Reduction of actomyosin contractility by treatment with fasudil or blebbistatin resulted in the disappearance of actin stress fibers observed in untreated cells (**a**) and the accumulation of actin signal in the cell cortex, as marked by arrowheads. **c**, Activation of Rho-kinase by leptin treatment increased the formation of actin stress fibers. **d**, Quantification of the actin signal in the peripheral regions of the cell versus the center demonstrates that fasudil treatment causes a greater than two-fold increase in actin deposition in the cell cortex. Leptin treatment, on the other hand, reduces this ratio significantly. **e-f**, Reduction of actomyosin contractility by fasudil treatment resulted in the peripheral distribution of pERM (red) signal along with actin (green). **g**, Membrane tether forces quantified by optical tweezers experiments significantly reduced upon leptin treatment. **h**, Reduced membrane tension by leptin treatment gave rise to significantly higher clathrin-mediated endocytosis dynamics. ***p<0.001, *p<0.05; two-tailed t test. Scale bar, 20 µm.

**Supplementary Figure 2.**
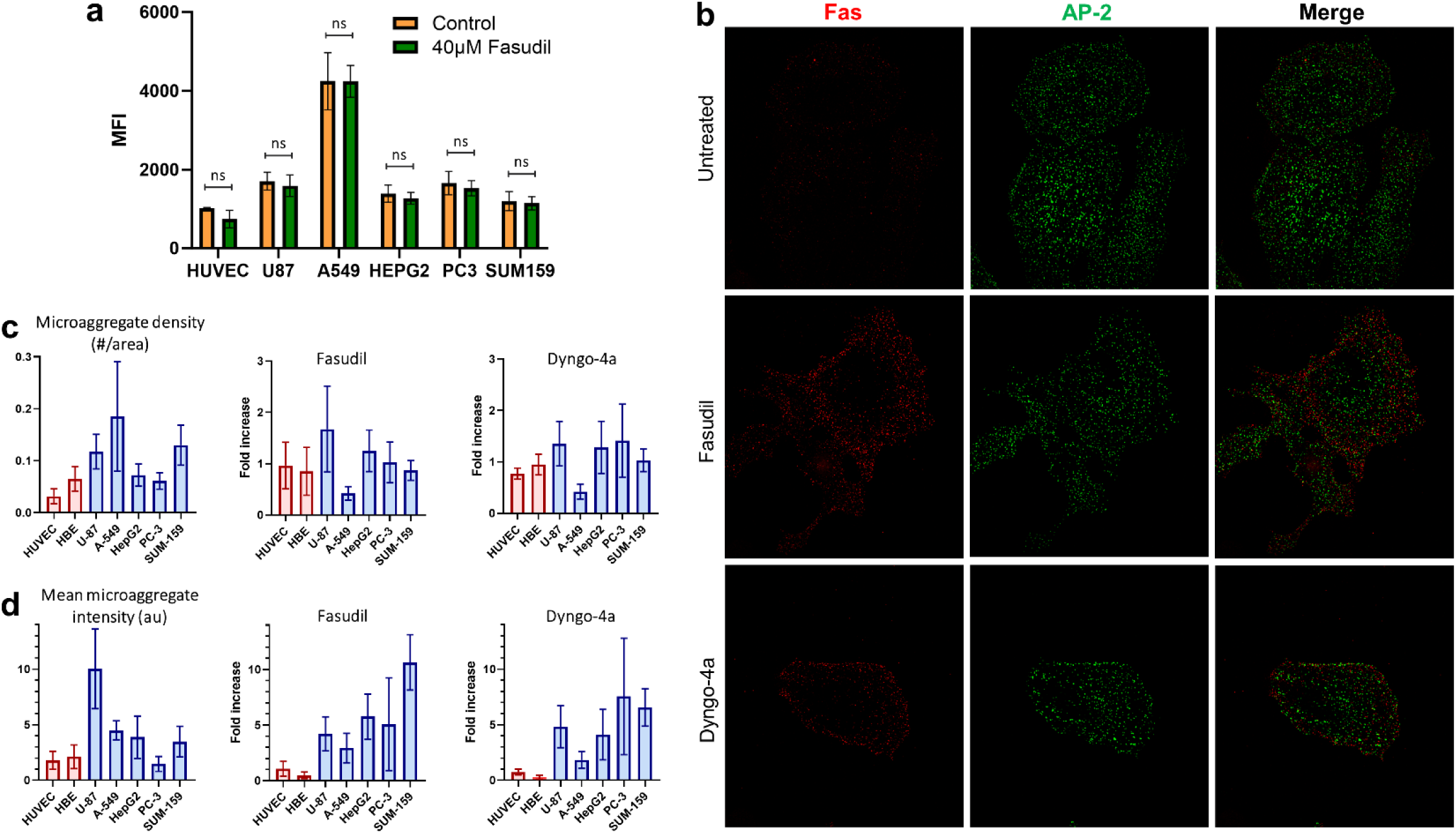
**a**, Flow cytometry analyses were conducted to test the effect of 40 µM fasudil on the total expression levels of Fas in HUVEC, U87, A549, HepG2, PC3 and SUM159 cells. The results are shown as mean fluorescence intensity +/− standard deviation. ns (nonsignificant), two-tailed t test. **b**, Spinning disk confocal fluorescence microscopy was used to image Fas microaggregates exclusively at the ventral/bottom surface of SUM159 cells that were genome edited to express AP2-EGFP, a marker of endocytic clathrin coats at the plasma membrane. The AP2-EGFP signal was used to verify that the image plane coincides with the ventral surface of the plasma membrane in these assays. **c**, The density of Fas microaggregates (mean ± sd) is shown for different cells (left). Treatment with 40 µM fasudil (middle) or 5 µM dyngo-4a (right) for 2 hours did not significantly increase microaggregate density in any of the cell types. **d**, Mean fluorescence intensity of Fas microaggregates (mean ± stdev) for different cells (left). Treatment with 40 µM fasudil (middle) or 5 µM dyngo-4a (right) for 2 hours did not increase the intensity of Fas microaggregates in HUVECs or HBE cells but did substantially increase this intensity in cancer cells (U87, A549, HepG2, PC3 and SUM159 cells). Scale bar, 20 µm.

**Supplementary Figure 3.**
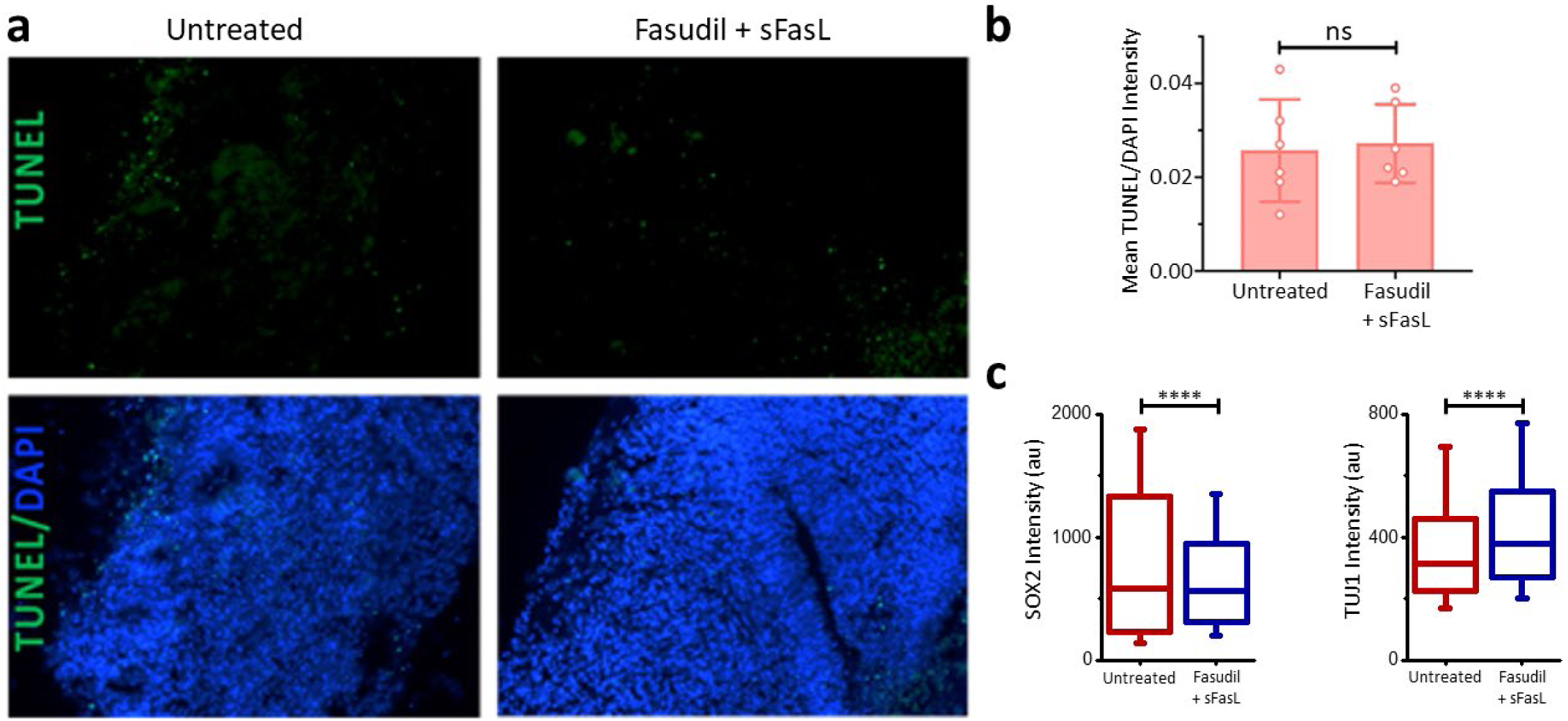
**a**, Seventy-eight-day-old healthy organoids were treated without or with both fasudil and sFasL. **b**, The ratio of TUNEL+ cells to the total number of nuclei was not significantly different in the combination treatment group and the untreated group (p = 0.79; unpaired t test). **c**, Box plots show the SOX2 and TUJ1 fluorescence staining levels in organoid cells before and after fasudil+sFasL treatment (N_CellsBefore_ = 1086; N_CellsAfter_ = 1506). We detected ∼13% increase in TUJ1 intensity (391.4 ± 9.0 vs 440.9 ± 6.1, mean ± sem) and ∼20% decrease in SOX2 intensity (833.3 ± 21.7 vs 695.8 ± 12.9, mean ± sem) upon treatment.

**Supplementary Figure 4.**
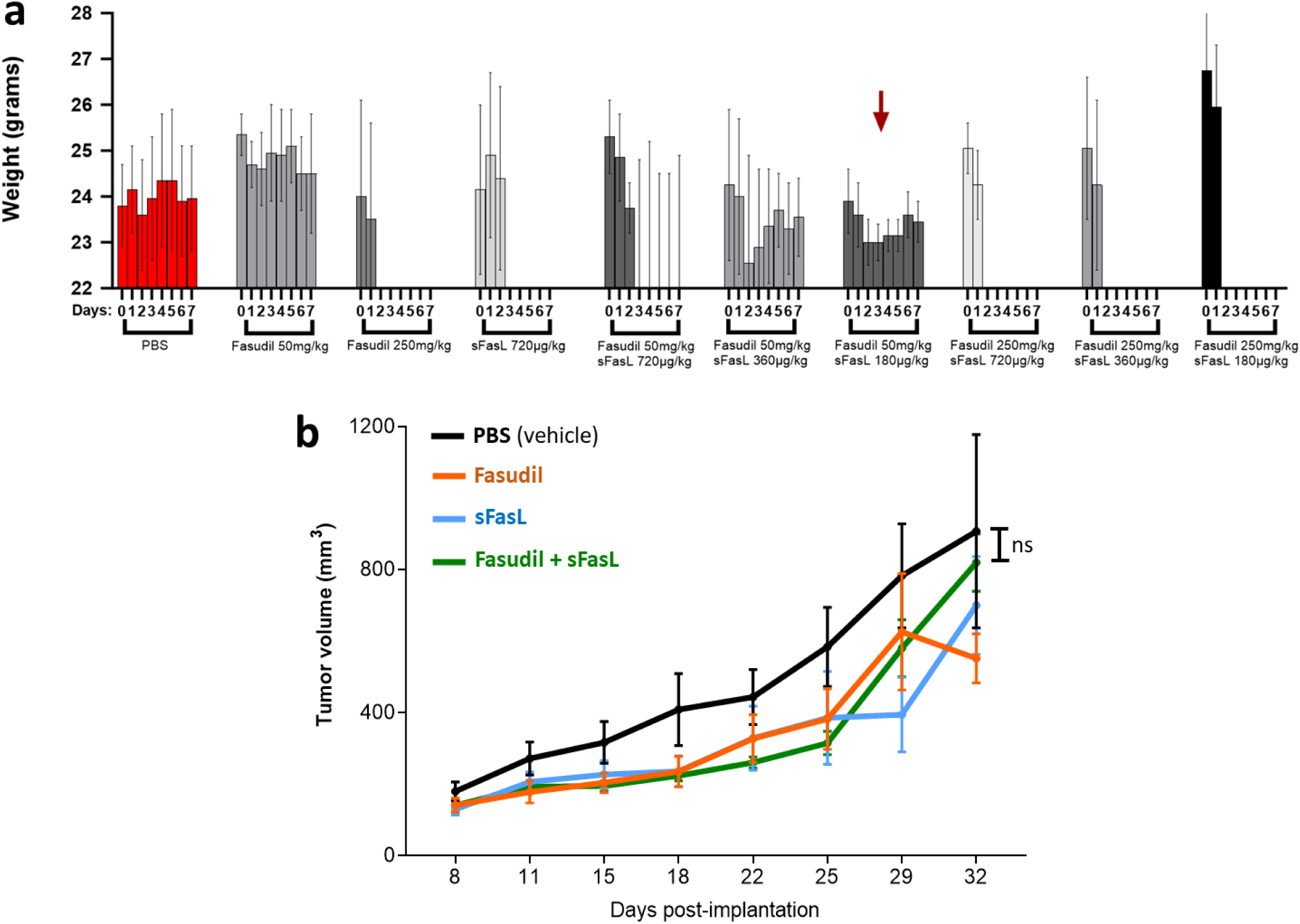
**a**, Results of the maximum tolerated dose assays show the weight of mice in different days of the intraperitoneal delivery of fasudil and/or sFasL at different concentrations. The doses used in our xenograft assays (50mg/kg fasudil and 180µg/kg sFasL) are marked with the red arrow. **b**, Intraperitoneal delivery of different treatments did not result in a significant difference in tumor volume (Ordinary one-way ANOVA). Overall, the fasudil+sFasL combination performed better than PBS, but the difference was not statistically significant.

## Supplementary Videos

**Video 1:** Z-stack acquired by spinning disk confocal imaging shows the surface localization of Fas (red) in a SUM159 cell genome edited to express AP2-EGFP (green).

**Video 2:** Untreated iPS-cardiomyocyte monolayer that is spontaneously contracting on the tissue culture plastic.

**Video 3:** 48 hours of treatment with combinations of fasudil – sFasL combination did not affect the contraction ability of iPS-cardiomyocyte monolayer.

